# When visual attention is divided in the flash-lag effect

**DOI:** 10.1101/2024.03.14.585092

**Authors:** Jane Yook, Hinze Hogendoorn, Gereon R. Fink, Simone Vossel, Ralph Weidner

## Abstract

The flash-lag effect (FLE) occurs when a flash’s position appears delayed relative to a continuously moving object, even though both are physically aligned. While several studies have demonstrated that reduced attention increases FLE magnitude, the precise mechanism underlying these attention-dependent effects remains elusive. In this study, we investigated the influence of visual attention on the FLE by manipulating the level of attention allocated to multiple stimuli moving simultaneously in different locations. Participants were cued to either focus on one moving stimulus or split their attention among two, three, or four moving stimuli presented in different quadrants. We measured trial-wise FLE to explore potential changes in the magnitude of perceived displacement and its trial-to-trial variability under different attention conditions. Our results reveal that FLE magnitudes were significantly larger when attention was divided among multiple stimuli compared to when attention was focused on a single stimulus, suggesting that divided attention considerably augments the perceptual illusion. However, FLE variability, measured as the coefficient of variation, did not differ between conditions, indicating that the consistency of the illusion is unaffected by divided attention. We discuss the interpretations and implications of our findings in the context of widely accepted explanations of the FLE within a dynamic environment.

## 1. Introduction

Attention is crucial in how we process and represent information in our visual environment. For dynamic information, attention may be essential for updating and maintaining coherent representations of moving objects (Iordanescu et al., 2009; Kerzel, 2003). When attention is limited, however, perceptual biases and illusions, such as the displacement in an object’s position due to its motion or the motion of other objects, can be significantly altered.

One such illusion is the flash-lag effect (FLE). The FLE occurs when a static flash appears next to a continuously moving object, leading to a perceived spatial offset between their positions despite their physical alignment at the moment of the flash (Nijhawan, 1994). Specifically, the flash is misperceived as lagging behind the moving object. Although several studies have demonstrated that the allocation of attention modulates the magnitude of the FLE, the precise mechanism underlying such attention-dependent effects remains elusive. Additionally, the relationship between attention and (in)variability of moving objects’ perceived positions is not well understood.

Sarich et al. (2007) employed a dual-task paradigm to compare the magnitude of the FLE in a flash-lag task performed alone or concurrently with a target detection task. They found that the FLE, measured as a point of subjective equality, was smallest when the flash-lag task was performed separately or with a slight interval from the detection task. Notably, when another task required simultaneous attention, the magnitude of the FLE increased, and detection worsened. This complements observations regarding the phenomenon of representational momentum (Freyd & Finke, 1984), where the perceived final position of a moving object is shifted forward in the direction of anticipated motion and increases with divided attention (Hayes and Freyd, 2002).

Shioiri et al. (2010) manipulated attention by varying the number of moving stimuli and presenting the flash next to one of these stimuli. They observed an increase in the FLE when attention was divided among two or six stimuli and a decrease when attention was focused solely on one stimulus. Conversely, when participants were pre-cued to the upcoming locations of either the moving object or the flash, the FLE reduced following valid cues compared to invalid (Baldo & Klein, 2010; Namba & Baldo, 2004; Vreven & Verghese, 2005; but see Khurana et al., 2000) or no cues (Shioiri et al., 2010). This pattern was also reported regarding representational momentum (Hubbard et al., 2009).

However, in tasks involving multiple object tracking (Pylyshyn & Storm, 1988), which requires precise localization of targets among other moving objects, divided attention has been shown to increase errors in extrapolating predictable motion trajectories (Adamian & Andersen, 2022; Howe & Holcombe, 2012; Luu & Howe, 2015; Zhong et al., 2014). These varying effects of reduced attention across different illusions and paradigms underscore the complexity of how attention influences the representation of moving objects and the need to disentangle the specific mechanisms underlying these effects. Therefore, the present study investigated whether attention not only influences the strength of perceptual illusions like the FLE but also affects the quality of representation, such as spatial resolution (Anton-Erxleben & Carrasco, 2013; Barbot & Carrasco, 2017; Yeshurun & Carrasco, 1998, 1999).

To this end, we examined whether divided attention affects the magnitude or the trial-to-trial variability of the FLE using trial-wise spatial cues. Previous studies often used stimuli grouped as a single moving object (e.g., dots arranged in a line; Baldo & Klein, 1995; Krekelberg & Lappe, 1999) or multiple objects following the same motion trajectory (e.g., dots revolving in a circular path; Khurana et al., 2000; Shioiri et al., 2010), which could introduce grouping effects. In contrast, our study employed objects characterized by independent motion trajectories to minimize such effects.

In our novel experimental paradigm, participants viewed an array of identically appearing bars presented in different locations. A cue preceding each trial indicated which bar(s) participants should track. While all bars rotated simultaneously, participants were required to covertly track the moving bar in the cued quadrant(s) of the display. Importantly, the perceptual configuration remained the same whether attention was focused on a single bar or divided across two, three, or four bars. During each trial, a target flash was presented next to one of the bars, and participants were instructed to indicate the position of the corresponding bar at the time of the flash. We compared the effects of divided attention to focused attention on both FLE magnitude and consistency (trial-to-trial variability of the FLE).

## 2. Materials and Methods

### 2.1 Participants

Twenty-six healthy adults were recruited from the Forschungszentrum Jülich to participate in the experiment. Two participants were excluded due to insufficient data quality (see section 2.5 for details), resulting in data from 24 participants (15 female, 9 male, age [M: 28.8 years, SD: 4 years], range 21–40 years) being included in the final analysis. The sample size was determined by a preliminary power calculation for a desired medium effect size (Cohen’s *f* = 0.25) with a power of 80% and an alpha level of 0.05 for a repeated-measures analysis of variance (ANOVA). All participants were fluent in spoken and written English, had normal or corrected-to-normal visual acuity, and had no neurological or psychiatric disorders. Handedness was not a selection criterion; participants self-reported their handedness preferences (19 right-handed, 5 left-handed). All participants provided informed written consent and received compensation of 10 Euros per hour for their participation. This study was conducted in accordance with the Declaration of Helsinki and approved by the ethics committee of the German Psychological Society (DGPs) (Ethics ID: 2022-02-03VA).

### 2.2 Apparatus

Visual stimuli were generated using Psychopy 3.0 (Peirce, 2007, 2009) and presented on a 22-inch Samsung SyncMaster 2233RZ LCD monitor with a resolution of 1680 x 1050 pixels and a refresh rate of 120 Hz (as outlined in Wang & Nikolic, 2011). A chinrest was used to stabilize the head and maintain a viewing distance of 70 cm, with the center of the screen approximately at eye level. A standard QWERTY keyboard was positioned below the chinrest and not visible from the participants’ field of view, with the left hand on the space bar and the right hand on the arrow keys. The experiment was conducted in a soundproof, light-attenuated room.

### 2.3 Stimuli

A white fixation cross, subtending 0.5° x 0.5° of visual angle (dva), was continuously displayed at the center of a black background. The spatial cue consisted of one, two, three, or four yellow dots (radius = 0.15 dva) presented at the corners of the central fixation cross, indicating the quadrant(s) where the target flash could occur. Every possible combination of dots was realized (e.g., for two quadrants, two dots on the left, right, upper, lower, or diagonal quadrants).

The moving stimuli consisted of four gray bars, each rotating smoothly around the center of its respective quadrant at an angular velocity of 240°/s. Each bar measured 0.13 x 6.04 dva, with its inner end positioned at 9.98 dva from the central fixation. On one end of each bar was a small dot (radius = 0.13 dva) used to indicate the bar’s position relative to the target flash. The target was a red dot of the same size (radius = 0.13 dva) and was always positioned at 0.8 dva from the bar’s dot end.

### 2.4 Procedure

In each trial, participants were instructed to fixate on the central fixation cross for the duration of the stimulation sequence (**Figure 1A**). Attention cues were displayed for 600 ms, pointing to the quadrant(s) to be attended. These cues pointed to the quadrant but not the specific position within the quadrant where the target was likely to appear in the upcoming display.

**Figure 1.**
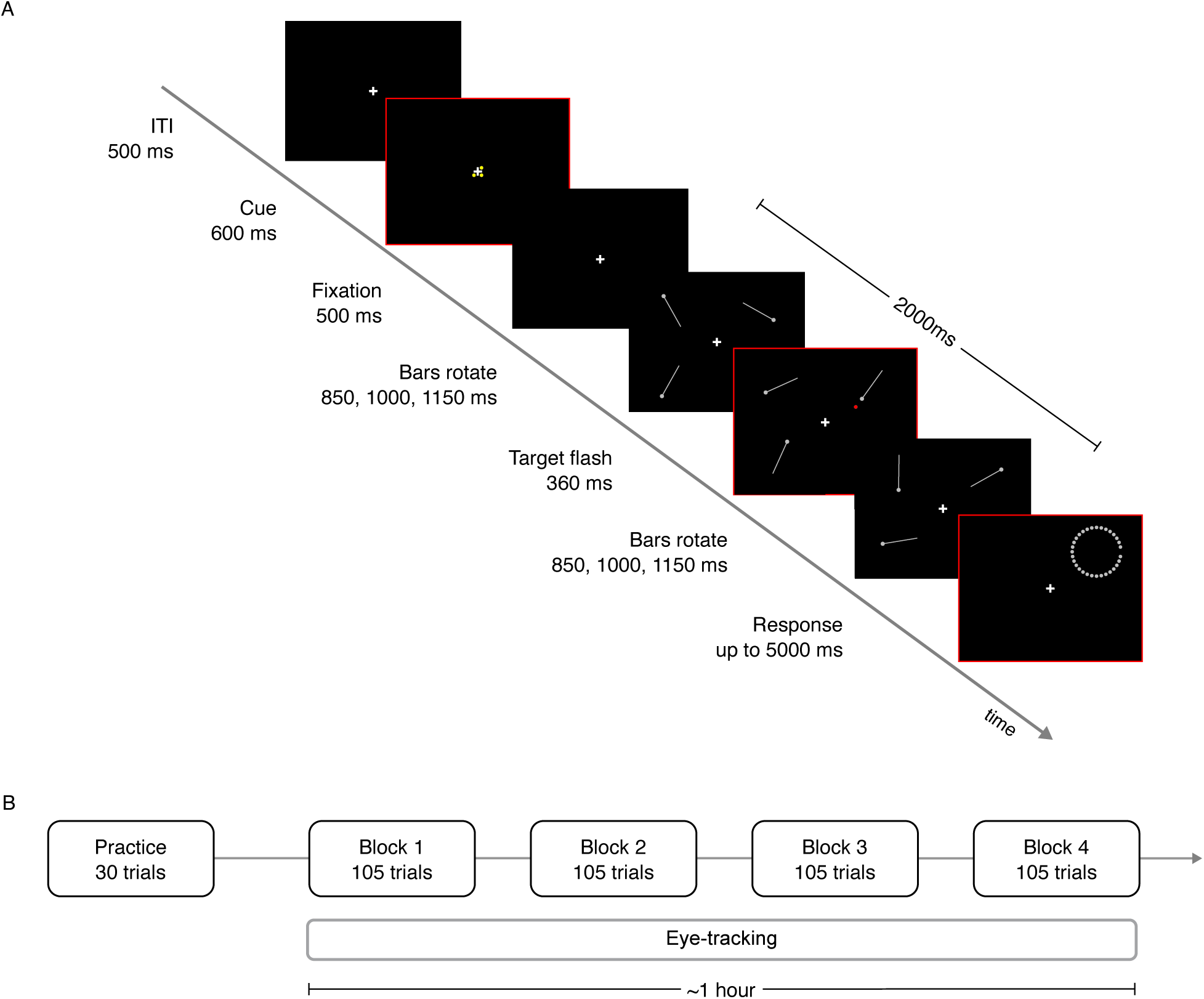
Experimental paradigm. (**A**) Trial sequence. Spatial attention cues were presented at the central fixation cross to indicate that the participants should orient their attention to the upcoming display, featuring an array of four rotating bars (not drawn to scale) in corresponding quadrants. The target flashed next to a bar after half the time of the overall sequence. Here, a target is presented at a cued quadrant, but sometimes the target appeared at a non-cued quadrant (see text for details). Following the sequence, a response ring was presented until participants made a response. The subsequent trial began after an inter-trial interval of 500 ms. Each trial lasted up to 8 s. (**B**) Procedure timeline. Participants performed at least 30 practice trials for task familiarization, followed by four blocks of 105 trials, totaling 420 trials over approximately one hour, including eye-tracking. Each block comprised trials from all four attention conditions and every combination of cue-target configurations with equal probability. The order of trials was randomized within the block.

After a 500-ms interval, allowing participants to covertly orient their attention to the cued quadrant(s), an array of four rotating bars was presented around the central fixation cross. The bars rotated in either a clockwise or counter-clockwise direction, alternating randomly across trials. Each bar started from a unique orientation, chosen from one of 16 uniformly distributed orientations. This configuration allowed the bars to rotate along independent motion trajectories despite their shared direction of motion.

Periodically after 850, 1000, or 1150 ms (chosen at random), the target briefly flashed (25 ms) next to one of the bars. After the target offset, the bars continued to rotate for an additional 1150, 1000, or 850 ms (for a total duration of 2000 ms). Participants’ task was to indicate the instantaneous position of the cued bar at the time of the flash on a circular response probe.

The response probe comprised a ring of 60 small grey dots (each dot radius = 0.13 dva) – whose diameter was equal to the length of a bar. To perform the task, participants navigated the ring using the left and right arrow keys to select their perceived position of the bar’s dot end displayed on the ring.

The starting position on the ring randomly varied from trial to trial, which introduced an inherent variability in how quickly participants could accurately respond, resulting in longer response times (RTs) when the distance from the desired position was greater. While RT is a common measure in similar paradigms (see Khurana et al., 2000), we did not analyze RTs for the current experiment. Instead, participants were instructed to respond within a 5000-ms window, focusing on accuracy rather than speed. A response was registered when the enter key was pressed. The subsequent trial began after an inter-trial interval (ITI) of 500 ms.

Importantly, since the target and its corresponding bar were presented in alignment, the perceived displacement of the stimulus positions in the direction of motion, or the lack thereof, directly determined the FLE for each trial. This method contrasts with the conventional approach of averaging offsets based on the point of subjective equality across blocks within a specific condition (cf. Yook et al., 2022).

The probe location corresponded to the quadrant where the target had been presented in the previous display, eliminating the need for participants to additionally report the target’s quadrant in each trial. However, the cue-target validity varied across trials. Specifically, the target appeared in the cued quadrant(s) 90% of the time. In the remaining 10% of trials, termed “catch trials”, the target appeared in a non-cued quadrant, and participants were instructed to press the space bar without indicating the bar’s position with the response probe. Catch trials ensured participants’ attention to the cues. Incorrect responses, such as responding using the probe in catch trials or pressing the space bar in non-catch trials, prompted feedback, “Please ATTEND TO THE CUES as they are helpful”, at fixation for 1000 ms. No feedback on behavioral performance was given otherwise.

The task was organized into four blocks, each consisting of 105 trials (96 non-catch trials and 9 catch trials; **Figure 1B**), totaling 420 trials conducted over approximately one hour. Each trial lasted up to 8 s. Every possible combination of cue-target configurations appeared within each block with equal probability. Participants underwent four attention conditions, and the order of trials was randomized within each block. To minimize fatigue, participants were instructed to take regular self-paced breaks between blocks and between every ∼5–10 mins within a given block, as indicated by a break screen.

Before the main experiment, participants completed at least 30 practice trials to familiarize themselves with the task. Upon completion of the experiment, participants were debriefed about the study’s purpose related to the FLE. None of the participants reported being aware that the positions of the attended bar and target were aligned.

### 2.5 Data quality assessment

Data from participants who failed to identify at least 75% of catch trials were excluded from the analysis to mitigate potential non-compliance with task instructions and task artifacts. This criterion ensured participants’ adherence to attending to the cued bar(s), as instructed. Data from two participants who detected an average of 2.5 catch trials (SD: 3.5) out of 36 throughout the experiment were excluded. Additionally, incorrectly responded non-catch trials were excluded. Among the remaining 24 participants, who on average detected 33.3 catch trials (SD: 2.4), data from an average of 379 correct non-catch trials (min: 371, max: 384, SD: 3.8) were available for analysis.

### 2.6 Behavioral analysis

Behavioral data were pre-processed using Python 3.5 (Van Rossum & Drake, 2009) within the Anaconda environment (Anaconda Inc., 2016) and analyzed in R Statistical Software v4.3.1 (R Core Team, 2023) with custom scripts.

During each trial, participants were instructed to direct their attention towards a single moving bar (focused attention condition) or spread their attention between two, three, or four bars (divided attention conditions: attend-to-two, -three, or -four). The FLE for each trial was determined by indexing the magnitude of the difference between the actual position of the target and the participant’s chosen position of the bar. A value of zero indicated no perceived difference between the positions of the two stimuli. Positive values indicated a flash-lag effect, where the moving bar was perceived as ahead of the flash. Conversely, negative values indicated a flash-lead effect, where the flash was perceived as ahead of the moving bar.

#### 2.6.1 Assessment of magnitude

The FLE values were aggregated to compute the median FLE magnitudes for each condition of each participant. We then compared these medians between the focused and divided attention conditions using a repeated-measures ANOVA using the four-level factor *attention*, with the ez package (Lawrence, 2016) in R. Following the ANOVA, we performed paired-sample *t-*tests to test the hypotheses that there would be significant differences in FLE magnitudes between focused attention and divided attention conditions, as well as between the various divided attention conditions To account for multiple comparisons, statistically significant differences were determined based on a threshold of 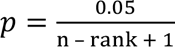 (one-tailed), adjusting for familywise error rate (FWE) using the Holm-Bonferroni correction (Holm, 1979). Greenhouse-Geisser corrections were applied when Mauchly’s test indicated a violation of sphericity.

To handle potential outliers, FLE values exceeding 3 standard deviations from the mean FLE values were identified and excluded. Analyses were performed both including and excluding outliers; however, no significant differences were observed when outliers were excluded. Therefore, the reported results include outliers, with descriptive statistics shown in **Appendix A, Table A1** to provide a more comprehensive insight into the distribution of the FLE values. To validate our findings against non-normality assumptions, we employed a Friedman’s test as a non-parametric alternative. The results of the Friedman’s test aligned with those of the repeated-measures ANOVA.

#### 2.6.2 Assessment of variability

In order to examine whether the distribution of FLE magnitudes became more variable with divided attention, the same procedure was repeated using the coefficient of variation (CV), calculated by dividing the standard deviation by the absolute mean of each condition of each participant. This allowed for comparing trial-to-trial variability across participants that vary widely in magnitude, by offering a relative measure of variability for comparing individuals.

### 2.7 Eye-tracking recording and analysis

We monitored eye movements using an infrared EyeLink 1000 Plus (SR Research Ltd., Mississauga, Canada) system at a sampling rate of 1000 Hz to evaluate how well participants had maintained fixation during the stimulation sequence of each trial. At the beginning of the experiment, the eye tracker was calibrated with a 5-point calibration procedure to establish an accurate gaze position of the left eye. Acceptable calibration values had to meet the validation criterion of < 0.5 dva and maximum error of < 1.5 dva. Due to technical issues, eye-tracking data was not recorded for 12 participants, and two additional datasets were incomplete.

Eye-tracking data were pre-processed and analyzed in R using the eyelinker package (Barthelme, 2021) and custom scripts. Fixations and saccades were determined from the raw gaze position data using Eyelink’s default event parser. A fixation was defined as an event lasting at least 100 ms, allowing deviation of 1-dva radius from the central fixation cross. Saccades were identified using a velocity threshold of 30°/s and acceleration threshold of 8000°/s^2^. Eye-blink events were excluded.

Across attention conditions, we examined the proportion of fixation time participants spent on the central fixation cross relative to the overall time during each of three distinct phases of a trial, wherein participants were instructed to maintain fixation:

1. During the cue presentation (cue) lasting 600 ms,
2. Immediately after the cue offset (orient) lasting 500 ms, and
3. Before and after the target presentation (target), with a combined duration of 2000 ms.

Differences in the proportion of fixation between the different attention conditions were analyzed with a repeated-measures ANOVA using the four-level factor *attention*.

### 2.8 Data and code availability

All experiment and analysis scripts are publicly available at https://osf.io/5pn9s/.

## 3. Results

### 3.1 Fixation controlled under all attention conditions

The proportions of fixation time across various phases of the experiment (cue, orient, and target) are presented in **Table 1**.

**Table 1.** Proportion of fixation time across attention conditions.

| ID | Cue |  | Orient |  | Target |  |
| --- | --- | --- | --- | --- | --- | --- |
|  | *M* | *SD* | *M* | *SD* | *M* | *SD* |
| 1 | 0.93 | 0.3 | 0.95 | 0.2 | 0.97 | 0.2 |
| 2 | 0.93 | 0.3 | 0.94 | 0.2 | 0.96 | 0.2 |
| 3 | 0.99 | 0.1 | 0.99 | 0.1 | 0.98 | 0.1 |
| 4 | 0.95 | 0.2 | 0.92 | 0.3 | 0.97 | 0.2 |
| 5 | 0.91 | 0.3 | 0.96 | 0.2 | 0.98 | 0.2 |
| 6 | 0.96 | 0.2 | 0.96 | 0.2 | 0.98 | 0.1 |
| 7 | 0.95 | 0.2 | 0.95 | 0.2 | 0.97 | 0.2 |
| 8 | 0.96 | 0.2 | 0.98 | 0.1 | 0.99 | 0.1 |
| 9 | 0.87 | 0.3 | 0.91 | 0.3 | 0.98 | 0.1 |
| 10 | 0.98 | 0.2 | 0.99 | 0.1 | 0.99 | 0.1 |

*Note.* The fixation duration may be influenced by interruptions such as blinks, signal fallout, or saccades. These events were initially identified and removed ahead of the fixation control analysis.

We performed a repeated-measures ANOVA to compare the proportion of fixation between different attention conditions within each phase. There were no significant differences in fixation across cue (one-way ANOVA, *attention*: *F*(3, 27) = 0.7, *p* = 0.5), orient (one-way ANOVA, *attention*: *F*(3, 27) = 0.5, *p* = 0.7), or target phases (one-way ANOVA, *attention*: *F*(3, 27) = 0.4, *p* = 0.8). Overall, these results indicate that participants consistently maintained a high degree of central fixation, regardless of whether their attention was focused on a single quadrant or divided across multiple quadrants.

### 3.2 Attentional modulation of FLE magnitude, but not trial-to-trial variability

Participants reported the perceived position of a moving bar, which was compared to its physical position. This task assessed the flash-lag effect (FLE), where the difference between the physical and perceived (reported) position of the bar at the time of the flash indicates the magnitude of the illusion. We analyzed FLE magnitudes, calculated as the average of individual medians, across different attention conditions, as shown in **Figure 2A**.

**Figure 2.**
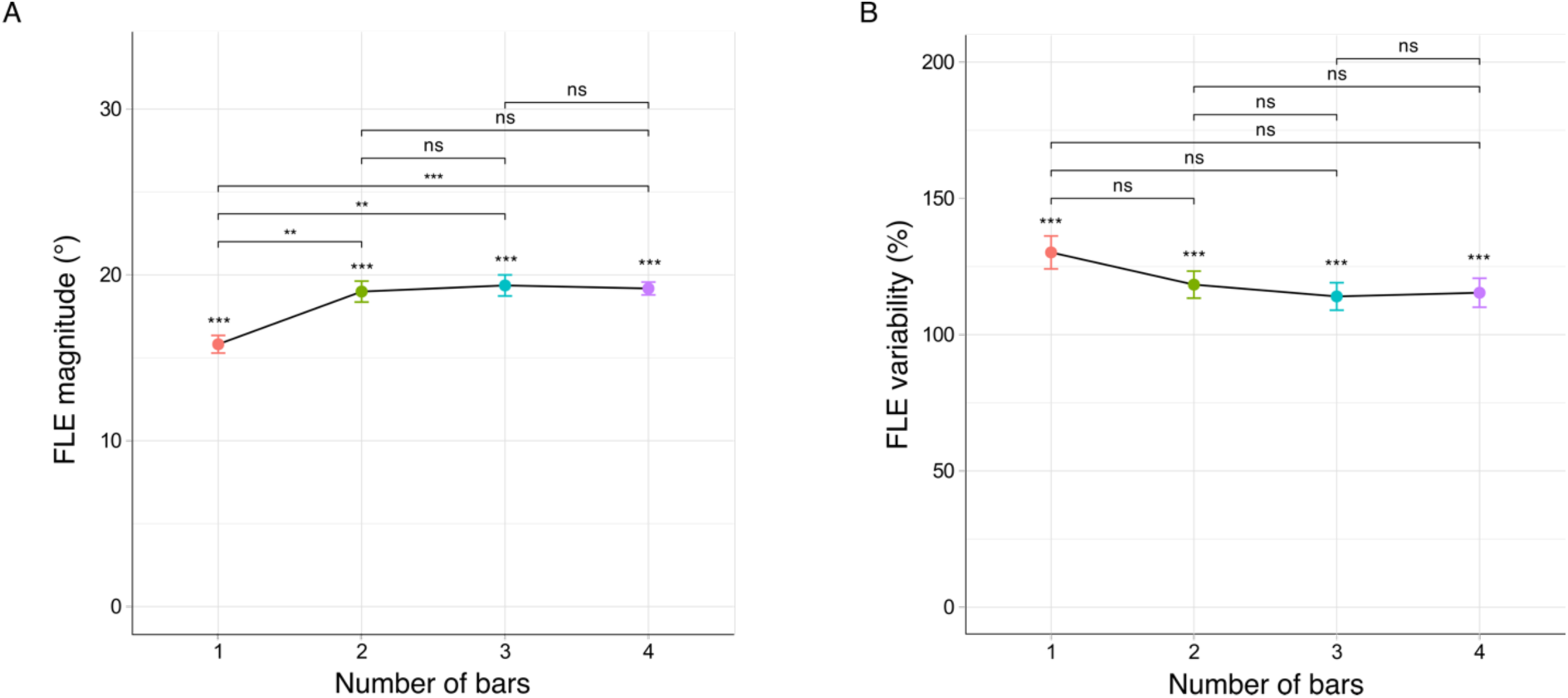
FLE results. (**A**) FLE magnitudes as a function of number of attended moving bars. All values are reported as means ± within-subject standard error of the mean (S.E.M., error bars). ns (not significant) denotes *p* > 0.05, ** denotes *p* <0.01, *** denotes *p* < 0.001. (**B**) The same described in A but for FLE variability (coefficient of variation).

Figure 2. FLE results. (**A**) FLE magnitudes as a function of number of attended moving bars. All values are reported as means ± within-subject standard error of the mean (S.E.M., error bars). ns (not significant) denotes *p* > 0.05, ** denotes *p* < 0.01, *** denotes *p* < 0.001. (**B**) The same described in **A** but for FLE variability (coefficient of variation).

The illusion was prominent across all attention conditions, averaging between 15-20° (see **Appendix A, Table A2**). Overall, one-way ANOVA revealed a significant effect of attention on the FLE magnitude (*F*(3, 69) = 9.2, *p* < 0.001). Specifically, the mean FLE magnitudes were significantly larger in divided attention conditions (two to four bars) compared to the focused attention condition (one bar), as detailed in **Table 2**.

**Table 2.**
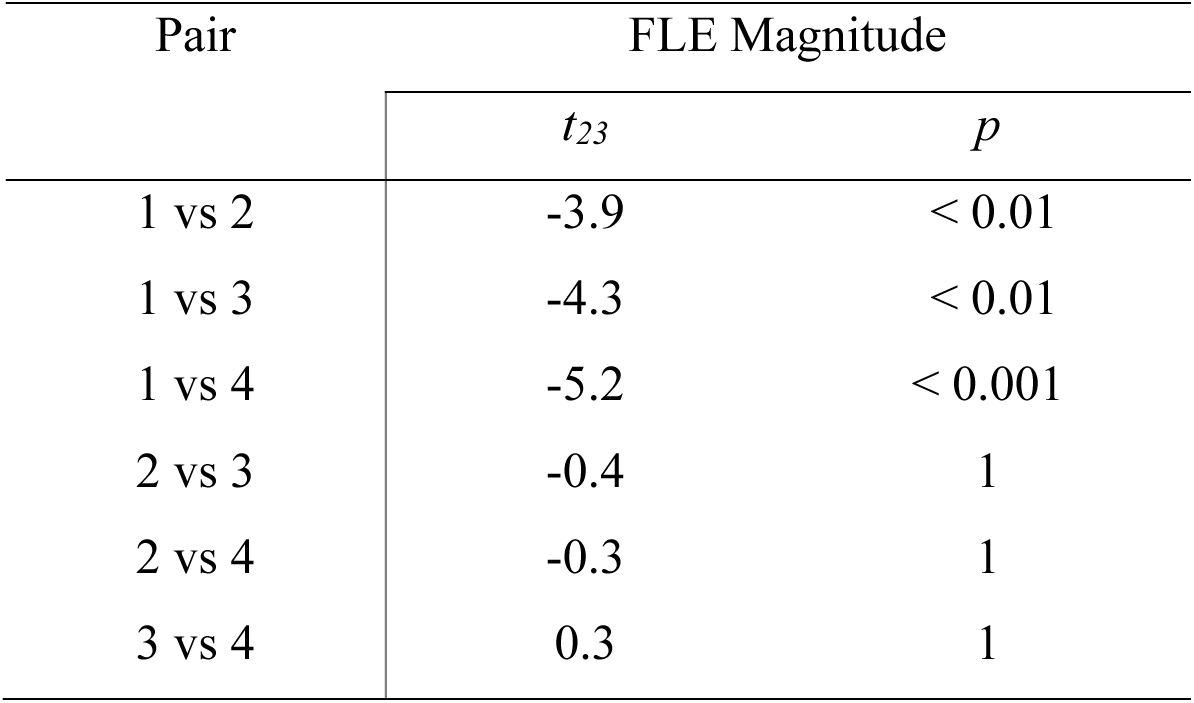
Paired-sample *t*-tests (one-tailed) results with correction for multiple comparisons for FLE magnitude.

Figure 2B (see also **Appendix B**) depicts a contrasting pattern in trial-to-trial FLE variability, as measured as the coefficient of variation (CV), with a larger CV in focused attention than in divided attention conditions. However, no significant differences emerged across conditions (one-way ANOVA, *attention*: *F*(3, 69) = 1.9, *p* = 0.1). Notably, the CVs exceeding 100% indicate a high degree of variability of FLE values relative to their mean, suggesting that the FLE values varied widely within participants (see also **Appendix A, Table A1**).

## 4. Discussion

In this study, we aimed to investigate how divided attention affects the perceived position of a moving object compared to a physically aligned static flash within the context of the FLE. The FLE is a well-documented illusion where a flash, when presented next to a moving object, appears to lag behind the moving object despite both being physically aligned. We found that FLE magnitude was augmented under divided attention conditions compared to focused attention. However, FLE variability across trials did not differ significantly between any of the attention conditions.

Participants viewed an array of four independently moving objects in distinct quadrants of the display during each trial. Attention was manipulated using spatial cues that directed participants to either focus on a single quadrant or to distribute attention across two, three, or four quadrants. This design allowed us to systematically compare FLE magnitude and trial-to-trial variability under different attentional loads. To ensure accurate allocation of attention, we included catch trials to verify that participants adhered to the provided cues. Subsequently, excluding catch responses from the analysis was crucial for differentiating between genuine attentional effects and potential confounding effects from incorrect cue compliance (see details below).

Our results revealed a clear effect of attention on the magnitude of the FLE. When attention was divided among multiple moving objects in different quadrants, FLE magnitudes increased significantly compared to when attention was focused on a single object. This finding aligns with previous studies (Namba & Baldo, 2004; Sarich et al., 2007; Shioiri et al., 2010) and is strengthened by our larger sample size (N = 24 compared to 15, 14, and 5, respectively) and a trial-wise measure of the FLE, which provides a more precise and nuanced understanding of the illusory effect. Interestingly, FLE variability did not show significant differences across the various attention conditions. These results suggest that while divided attention in the FLE amplifies the FLE magnitude, it does not necessarily lead to greater variability in representations from trial to trial compared to focused attention. However, it is important to note that the individual distributions of the FLE values contributed to large coefficients of variation (CV).

Moreover, the increase in FLE magnitude was significant only when shifting from one object to multiple objects, with no further increases when attention was distributed among two, three, or four objects. This pattern is consistent with Shioiri et al.’s (2010) findings, which involved up to six dots arranged in a circle. By contrast, Hogendoorn et al. (2010) observed a gradual increase in FLE magnitude, measured as time, with a step-wise increase in FLE variability from one to two clocks, with no further increases beyond that. This suggests a saturation effect at the condition where attention was focused on two objects, indicating that further dividing attention did not significantly impact the FLE under highly divided-attention conditions (Howe & Holcombe, 2012; Luu & Howe, 2015).

Two possible explanations arise from these observations. Firstly, as proposed by Shiori et al. (2010), the saturation effect may relate more to the spatial spread of attention than to the number of attended objects. For instance, Khurana et al. (2000) observed no changes in FLE magnitudes when one of two possible flash locations (either above or below the fixation point) was cued, similar to our study. This suggests that the effect of attention may not follow a straightforward pattern but is highly dependent on task complexity (Khurana et al., 2000; Shioiri et al., 2010). Secondly, attentional resources might be constrained by hemifield (Alvarez & Cavanagh, 2005; Luck et al., 1989) or even by quadrants (Carlson et al., 2007, 2011). If each quadrant has a limited attentional capacity, further dividing attention might not affect the representation of additional moving objects in extra quadrants. This could account for the deviations noted in our analysis of anisotropies (see **Appendix C**) and the discrepancies observed in Khurana et al. (2000), where no additional attentional modulation was found compared to other FLE studies considered here. Future research could explore these mechanisms of attentional saturation by manipulating perceptual load, such as varying the number of stimuli (> 1) within each quadrant or adjusting their eccentricities (Baldo et al., 2002). Additionally, exploring the impact of motion speed (Shioiri et al., 2010) could further provide insights into whether increased attention is needed to track faster motion (cf. Yook et al., 2022), potentially influencing FLE magnitude to a greater extent than observed here.

Our findings align with several prominent theoretical accounts of the FLE, including theories of sequential processing including temporal integration (Krekelberg & Lappe, 1999, 2000), discrete sampling perception (Schneider, 2018), attention shifting (Baldo & Klein, 1995), and postdiction (Brenner & Smeets, 2000; Eagleman & Sejnowski, 2000). For specific details of each of these theories, readers are referred to Hogendoorn (2020), Holcombe and Corbett (2023), Hubbard (2014), and Schneider (2018). Arguably, when attention is divided, the brain has fewer resources to process each object’s motion trajectory, reducing information processing speed. This may contribute to increased delays in the visual system’s ability to extract relevant motion and position information for each object, as the brain needs to switch between multiple objects, including the flash. This delay in processing time results in a more considerable latency difference between the moving objects and the flash, augmenting the FLE.

Under the predictive motion explanation theory (Nijhawan, 1994), these compounded delays across multiple objects may necessitate more compensation, thereby increasing the FLE magnitude. The brain might overcompensate if it prioritizes efficiency over accuracy when processing multiple moving objects under dynamic conditions, such as in this experiment. In principle, this would allow the brain to simultaneously monitor and update representations of multiple objects, even when sensory information is delayed. According to this interpretation, the attention-dependent effects of perceptual biases and illusions outlined in the **Introduction** can be reconciled. When attention is reduced in representational momentum and the FLE, the visual system may rely more heavily on existing predictions rather than on slowly-arriving sensory input, enhancing the perceived forward displacement in moving objects. In contrast, reduced attention results in decreased extrapolation during multiple object tracking. This occurs because the task requires continuous and precise tracking of several objects, and the brain cannot allocate enough resources to accurately predict each object’s motion trajectory, leading to less effective extrapolation. Hence, the impact of reduced or divided attention varies depending on the nature of the task. While our findings are open to interpretation, we believe our findings contribute to this body of evidence by revealing the role of attention in efficiently processing multiple moving objects in dynamic environments. By contrast, our findings do not seem to align with the differential processing latencies theory (Whitney et al., 2000; Whitney & Murakami, 1998), which would predict that as moving objects are processed faster than static objects, divided attention would lead to a smaller processing latency difference between the two and a smaller FLE magnitude.

Taken together, these results suggest that, in the context of the FLE, attention primarily influences the rate at which events are processed within the visual system, rather than the quality of processing. While overall perception is certainly impacted by the quality of representation, the FLE appears to be more a result of how the visual system effectively manages highly attention-demanding and dynamic environments. This suggests that when attentional resources are divided among multiple moving objects, delays in accessing relevant information may accumulate (Carrasco & McElree, 2001; Giordano et al., 2009), leading to slower recognition of motion and delayed availability of position information for each object.

An alternative interpretation is that attention affects the anticipatory processing of the flash (Baldo et al., 2002; Vreven & Verghese, 2005), potentially reducing the latency difference between the flash and the moving object when attention is focused. Although it is challenging to rule out an effect of divided attention on the flash, this alone is unlikely to account for our results. This interpretation would also predict increased FLE magnitudes with divided attention, but it assumes that participants might anticipate the flash even without explicit cues. Previous studies (Baldo et al., 2002; Sarich et al., 2007; Shioiri et al., 2010; Vreven & Verghese, 2005) may have been confounded by this. However, our study’s use of catch trials reduces the likelihood that participants were simply waiting for the flash. If participants had been passively anticipating the flash, then they would have reported the FLE regardless of the attentional cues. Our data show that participants did notice the flash when it appeared next to a non-cued bar, suggesting that they did not merely wait for the flash on every trial. While our analyses concentrated on correctly responded non-catch trials, future research could include a separate condition for catch trials. A lack of discernable difference in FLE magnitudes between catch and non-catch conditions would strongly suggest that the changes observed were primarily driven by the flash itself. This would offer further insights into how attention affects the representation of moving objects rather than the detection of the flash.

In conclusion, the experiment reported here is the first to manipulate attention in the flash-lag paradigm combining conventional attentional cueing with divided attention procedures, alongside trial-wise FLE readouts. Our results demonstrate that the magnitude of motion-induced illusory perceptual effects varies with the level of attention in dynamic environments.

## Acknowledgments

We thank our colleagues at the INM-3 for their valuable discussions and support.

## Commercial relationships

None

## Appendices for When visual attention is divided in the flash-lag effect

Jane Yook *et al*.

## Appendix A: Descriptive statistics of FLE magnitude

**Table A1.**
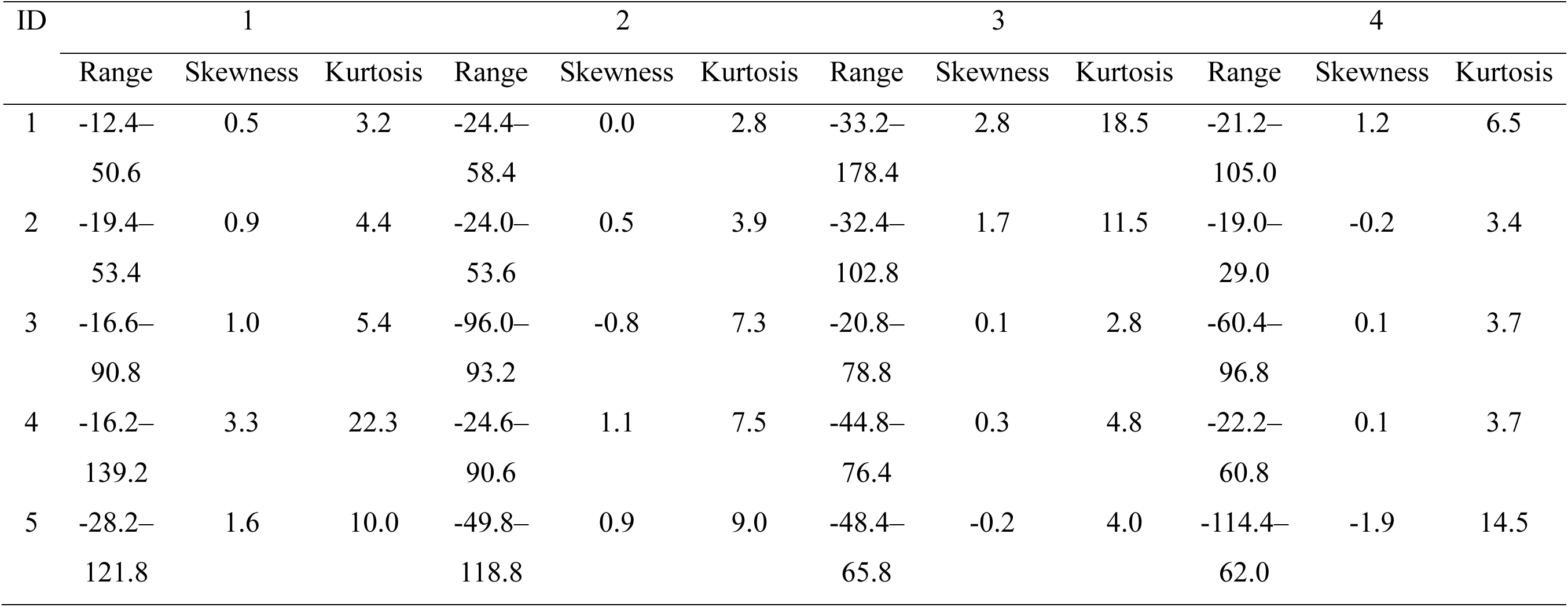
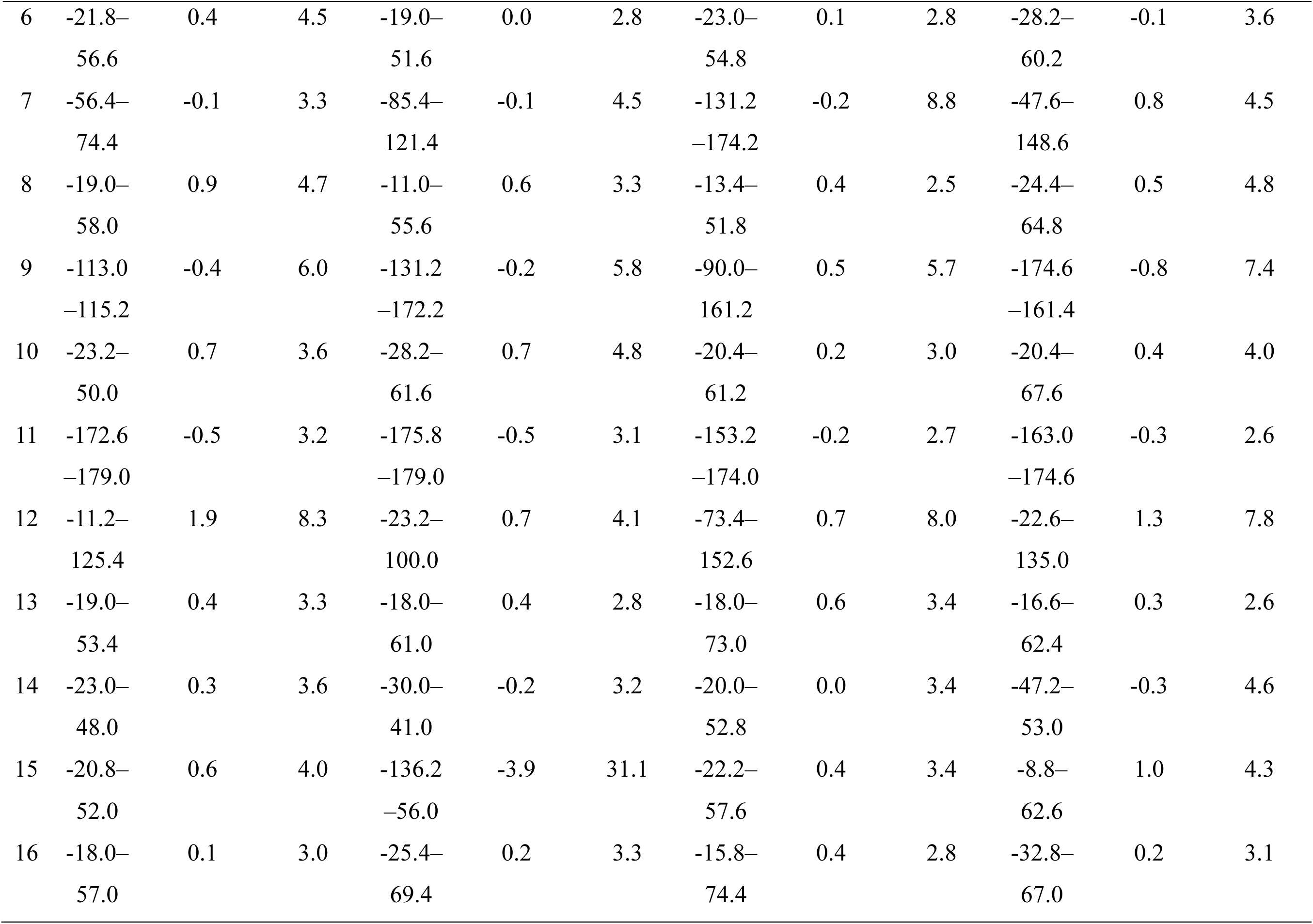
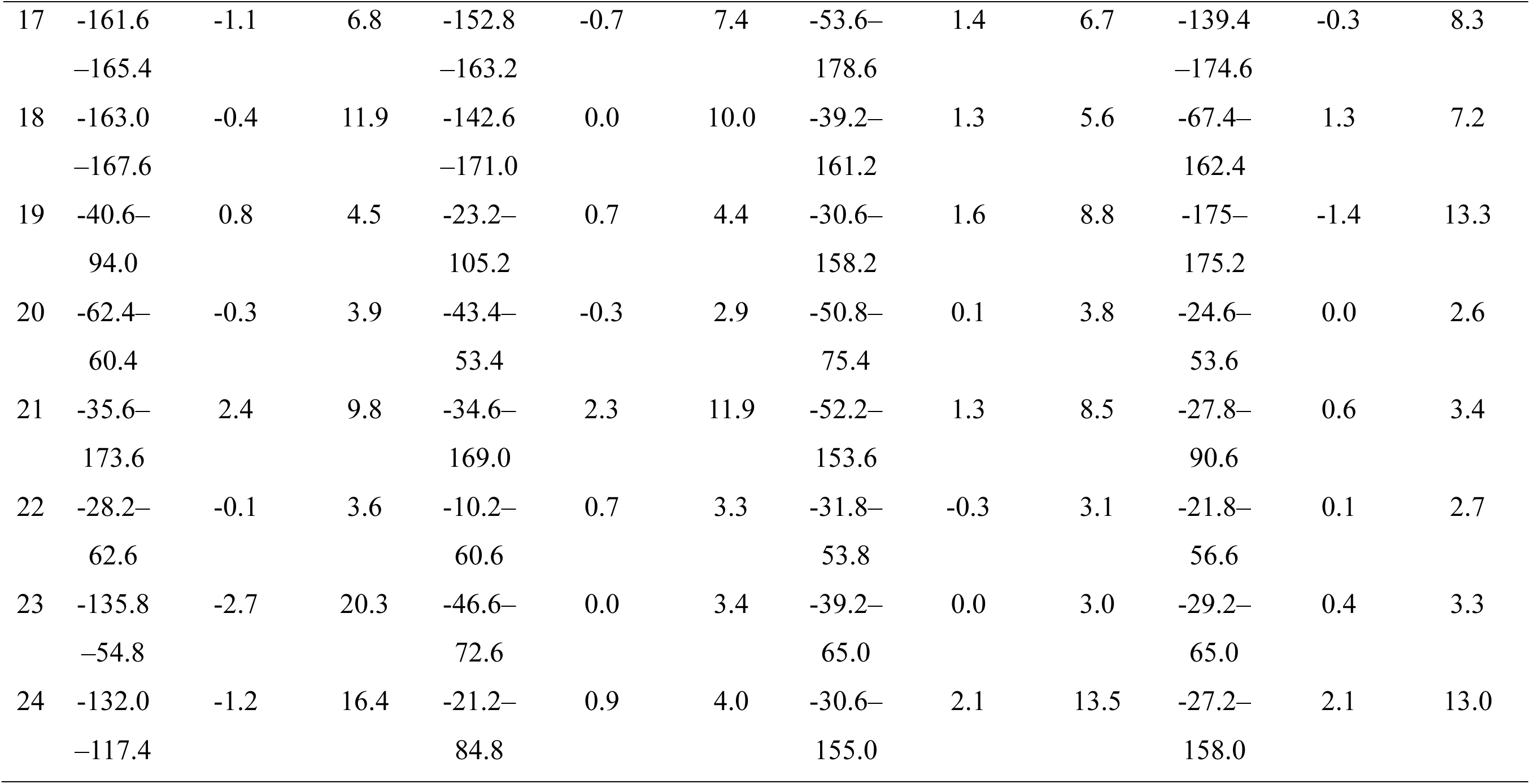
Descriptive statistics of FLE magnitudes across attention conditions.

**Table A2.**
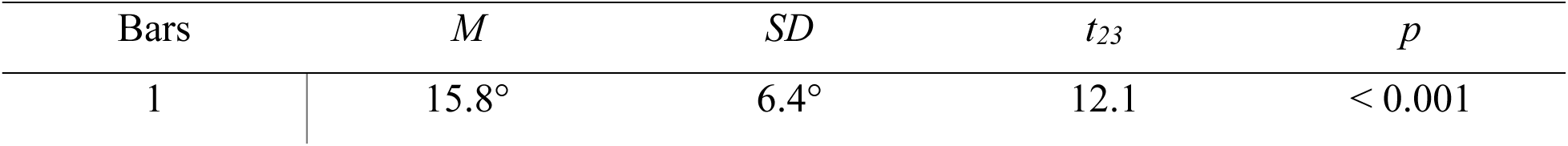
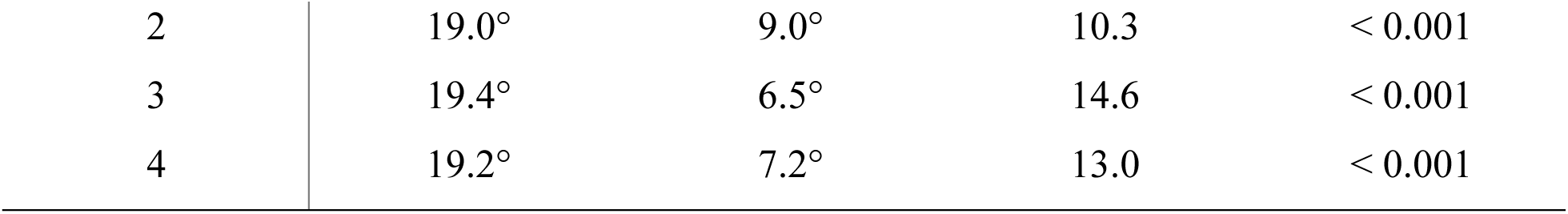
One-sample *t-*test (one-tailed) results against zero.

## Appendix B: Descriptive statistics of FLE variability

**Table B1.**
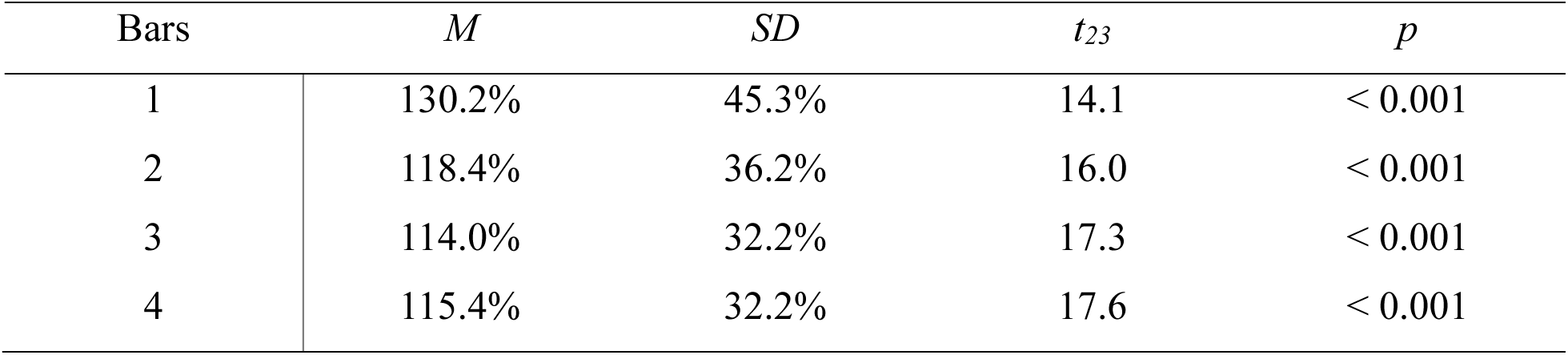
One-sample *t-*test (one-tailed) results against zero.

## Appendix C: Hemifield effects in the FLE

**Figure C1.**
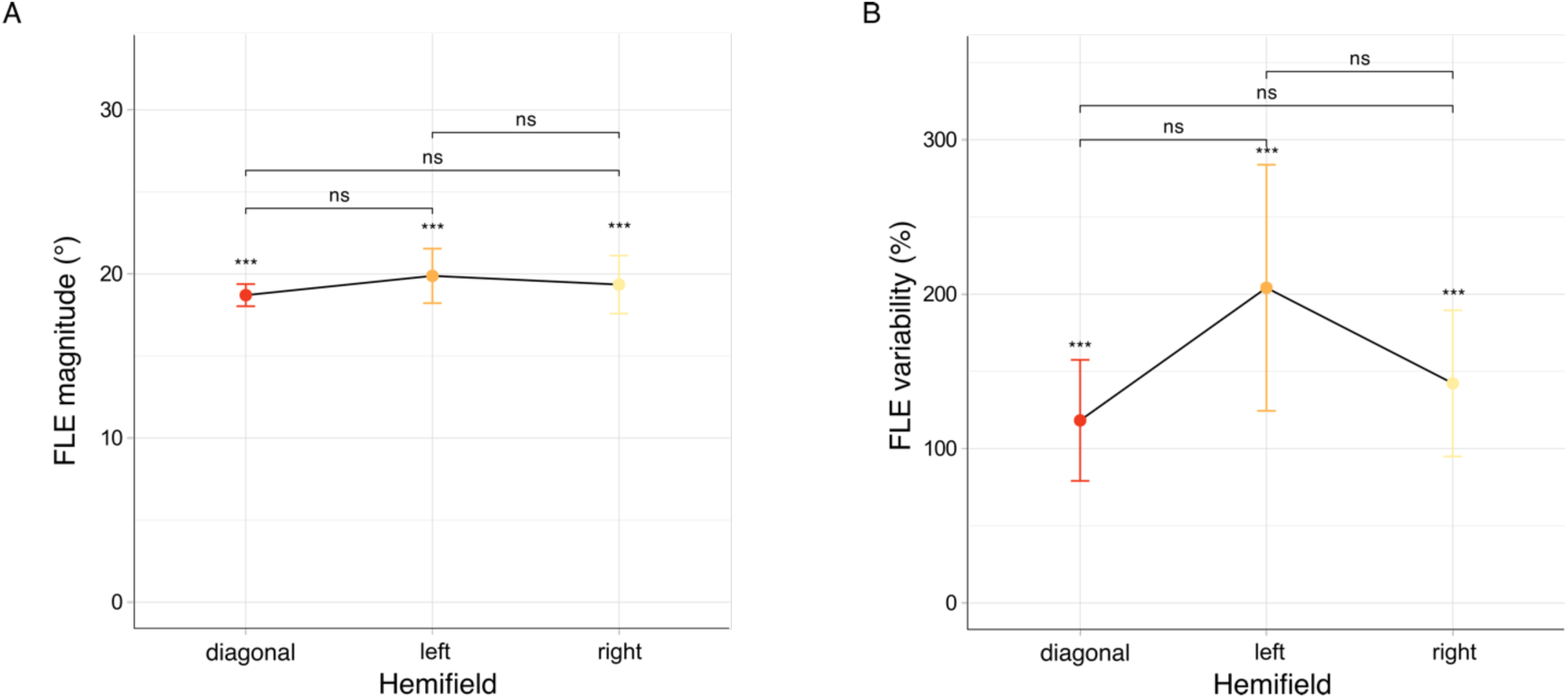
Hemifield results. Previous studies using linear motion have demonstrated anisotropies in the left and right sides of the visual field (Kanai et al., 2004; Shi & Nijhawan, 2008; Suzuki et al., 2023). Notably, the FLE tends to be more substantial when stimuli are presented in the left hemifield. Here, we aimed to test whether the effect of divided attention could be pronounced with these anisotropies, particularly within the attend-to-two condition. To achieve this, we examined the magnitude and variability of the FLE on trials where participants were cued to two quadrants within their left, right, or diagonally across the upper and lower visual hemifields. (**A**) The FLE magnitudes are plotted as a function of the to-be-attended hemifields. All values are reported as means ± within-subject standard error of the mean (S.E.M., error bars). ns (not significant) denotes *p* > 0.05. Although the FLE magnitude in the left hemifield was marginally higher than in both the right or diagonal hemifields, no significant modulation of attention was detected (one-way ANOVA, *hemiField: F*(2,46) = 0.2, *p* = 0.7). (**B**) The same as described in **A** but for FLE variability (one-way ANOVA, *hemiField: F*(2,46) = 0.6, *p* = 0.5).

## References

Adamian, N., & Andersen, S. K. (2022). Attentional Enhancement of Tracked Stimuli in Early Visual Cortex Has Limited Capacity. The Journal of Neuroscience, 42(46), 8709–8715. 10.1523/JNEUROSCI.0605-22.2022

Alvarez, G. A., & Cavanagh, P. (2005). Independent Resources for Attentional Tracking in the Left and Right Visual Hemifields. Psychological Science, 16(8), 637–643. 10.1111/j.1467-9280.2005.01587.x

Anaconda Software Distribution (Version 2-2.4.0). (2016). [Computer software]. Anaconda, Inc. https://www.anaconda.com/

Anton-Erxleben, K., & Carrasco, M. (2013). Attentional enhancement of spatial resolution: Linking behavioural and neurophysiological evidence. Nature Reviews. Neuroscience, 14(3), 188–200. 10.1038/nrn3443

Baldo, M. V. C., Kihara, A. H., Namba, J., & Klein, S. A. (2002). Evidence for an Attentional Component of the Perceptual Misalignment between Moving and Flashing Stimuli. Perception, 31(1), 17–30. 10.1068/p3302

Baldo, M. V. C., & Klein, S. A. (1995). Extrapolation or attention shift? Nature, 378(6557), Article 6557. 10.1038/378565a0

Baldo, M. V. C., & Klein, S. A. (2010). Paying attention to the flash-lag effect. In R. Nijhawan & B. Khurana (Eds.), Space and Time in Perception and Action (1st ed., pp. 396–407). Cambridge University Press. 10.1017/CBO9780511750540.023

Barbot, A., & Carrasco, M. (2017). Attention Modifies Spatial Resolution According to Task Demands. Psychological Science, 28(3), 285–296. 10.1177/0956797616679634

Barthelme, S. (2021). eyelinker: Import ASC Files from EyeLink Eye Trackers (Version 0.2.1) [Computer software]. https://cran.r-project.org/web/packages/eyelinker/index.html

Brenner, E., & Smeets, J. B. J. (2000). Motion extrapolation is not responsible for the flash–lag effect. Vision Research, 40(13), 1645–1648. 10.1016/S0042-6989(00)00067-5

Carlson, T. A., Alvarez, G. A., & Cavanagh, P. (2007). Quadrantic deficit reveals anatomical constraints on selection. Proceedings of the National Academy of Sciences, 104(33), 13496–13500. 10.1073/pnas.0702685104

Carlson, T. A., Cho, H., Turret, J., & Dakin, S. (2011). Psychoanatomy of visual attention: A unified account of quadrant and hemifield effects. Journal of Vision, 11(11), 101–101. 10.1167/11.11.101

Carrasco, M., & McElree, B. (2001). Covert attention accelerates the rate of visual information processing. Proceedings of the National Academy of Sciences, 98(9), 5363–5367. 10.1073/pnas.081074098

Eagleman, D. M., & Sejnowski, T. J. (2000). Motion Integration and Postdiction in Visual Awareness. Science, 287(5460), 2036–2038. 10.1126/science.287.5460.2036

Freyd, J. J., & Finke, R. A. (1984). Representational momentum. Journal of Experimental Psychology: Learning, Memory, and Cognition, 10(1), 126–132. 10.1037/0278-7393.10.1.126

Giordano, A. M., McElree, B., & Carrasco, M. (2009). On the automaticity and flexibility of covert attention: A speed-accuracy trade-off analysis. Journal of Vision, 9(3), 30. 10.1167/9.3.30

Hayes, A. E., & Freyd, J. J. (2002). Representational momentum when attention is divided. Visual Cognition, 9(1–2), 8–27. 10.1080/13506280143000296

Hogendoorn, H. (2020). Motion Extrapolation in Visual Processing: Lessons from 25 Years of Flash-Lag Debate. The Journal of Neuroscience, 40(30), 5698–5705. 10.1523/JNEUROSCI.0275-20.2020

Hogendoorn, H., Carlson, T. A., Vanrullen, R., & Verstraten, F. A. J. (2010). Timing divided attention. Attention, Perception, & Psychophysics, 72(8), 2059–2068. 10.3758/BF03196683

Holcombe, A. O., & Corbett, J. (2023). Temporal errors: Researchers should stop studying the flash-lag effect. 10.31234/osf.io/swzr7

Holm, S. (1979). A Simple Sequentially Rejective Multiple Test Procedure. Scandinavian Journal of Statistics, 6(2), 65–70.

Howe, P. D. L., & Holcombe, A. O. (2012). Motion information is sometimes used as an aid to the visual tracking of objects. Journal of Vision, 12(13), 10–10. 10.1167/12.13.10

Hubbard, T. L. (2014). The flash-lag effect and related mislocalizations: Findings, properties, and theories. Psychological Bulletin, 140(1), 308–338. 10.1037/a0032899

Hubbard, T. L., Kumar, A. M., & Carp, C. L. (2009). Effects of spatial cueing on representational momentum. Journal of Experimental Psychology: Learning, Memory, and Cognition, 35(3), 666–677. 10.1037/a0014870

Iordanescu, L., Grabowecky, M., & Suzuki, S. (2009). Demand-based dynamic distribution of attention and monitoring of velocities during multiple-object tracking. Journal of Vision, 9(4), 1–1. 10.1167/9.4.1

Kanai, R., Sheth, B. R., & Shimojo, S. (2004). Stopping the motion and sleuthing the flash-lag effect: Spatial uncertainty is the key to perceptual mislocalization. Vision Research, 44(22), 2605–2619. 10.1016/j.visres.2003.10.028

Kerzel, D. (2003). Attention maintains mental extrapolation of target position: Irrelevant distractors eliminate forward displacement after implied motion. Cognition, 88(1), 109–131. 10.1016/S0010-0277(03)00018-0

Khurana, B., Watanabe, K., & Nijhawan, R. (2000). The Role of Attention in Motion Extrapolation: Are Moving Objects ‘Corrected’ or Flashed Objects Attentionally Delayed? Perception, 29(6), 675–692. 10.1068/p3066

Krekelberg, B., & Lappe, M. (1999). Temporal recruitment along the trajectory of moving objects and the perception of position. Vision Research, 39(16), 2669–2679. 10.1016/S0042-6989(98)00287-9

Krekelberg, B., & Lappe, M. (2000). A model of the perceived relative positions of moving objects based upon a slow averaging process. Vision Research, 40(2), 201–215. 10.1016/S0042-6989(99)00168-6

Lawrence, M. (2016). ez: Easy Analysis and Visualization of Factorial Experiments (Version 4.4-0) [R]. https://CRAN.R-project.org/package=ez

Luck, S. J., Hillyard, S. A., Mangun, G. R., & Gazzaniga, M. S. (1989). Independent hemispheric attentional systems mediate visual search in split-brain patients. Nature, 342(6249), 543–545. 10.1038/342543a0

Luu, T., & Howe, P. D. L. (2015). Extrapolation occurs in multiple object tracking when eye movements are controlled. Attention, Perception, & Psychophysics, 77(6), 1919–1929. 10.3758/s13414-015-0891-8

Namba, J., & Baldo, M. V. C. (2004). The Modulation of the Flash-Lag Effect by Voluntary Attention. Perception, 33(5), 621–631. 10.1068/p5212

Nijhawan, R. (1994). Motion extrapolation in catching. Nature, 370(6487), 256–256. 10.1038/370256a0

Peirce, J. W. (2007). PsychoPy—Psychophysics software in Python. Journal of Neuroscience Methods, 162(1), 8–13. 10.1016/j.jneumeth.2006.11.017

Peirce, J. W. (2009). Generating stimuli for neuroscience using PsychoPy. Frontiers in Neuroinformatics, 2. https://www.frontiersin.org/articles/10.3389/neuro.11.010.2008

Pylyshyn, Z. W., & Storm, R. W. (1988). Tracking multiple independent targets: Evidence for a parallel tracking mechanism. Spatial Vision, 3(3), 179–197. 10.1163/156856888x00122

R Core Team. (2023). R: A Language and Environment for Statistical Computing (Version 4.3.1) [Computer software]. R Foundation for Statistical Computing. https://www.r-project.org/

Sarich, D., Chappell, M., & Burgess, C. (2007). Dividing attention in the flash-lag illusion. Vision Research, 47(4), 544–547. 10.1016/j.visres.2006.09.029

Schneider, K. A. (2018). The Flash-Lag, Fröhlich and Related Motion Illusions Are Natural Consequences of Discrete Sampling in the Visual System. Frontiers in Psychology, 9, 1227. 10.3389/fpsyg.2018.01227

Shi, Z., & Nijhawan, R. (2008). Behavioral significance of motion direction causes anisotropic flash-lag, flash-drag, flash-repulsion, and movement-mislocalization effects. Journal of Vision, 8(7), 24. 10.1167/8.7.24

Shioiri, S., Yamamoto, K., Oshida, H., Matsubara, K., & Yaguchi, H. (2010). Measuring attention using flash-lag effect. Journal of Vision, 10(10), 10–10. 10.1167/10.10.10

Suzuki, Y., Atmaca, S., & Laeng, B. (2023). The lateralized flash-lag illusion: A psychophysical and pupillometry study. Brain and Cognition, 166, 105956. 10.1016/j.bandc.2023.105956

Van Rossum, G., & Drake, F. L. (2009). Python 3 Reference Manual. CreateSpace.

Vreven, D., & Verghese, P. (2005). Predictability and the Dynamics of Position Processing in the Flash-Lag Effect. Perception, 34(1), 31–44. 10.1068/p5371

Wang, P., & Nikolic, D. (2011). An LCD Monitor with Sufficiently Precise Timing for Research in Vision. Frontiers in Human Neuroscience, 5. https://www.frontiersin.org/articles/10.3389/fnhum.2011.00085

Whitney, D., & Murakami, I. (1998). Latency difference, not spatial extrapolation. Nature Neuroscience, 1(8), 656–657. 10.1038/3659

Whitney, D., Murakami, I., & Cavanagh, P. (2000). Illusory spatial offset of a flash relative to a moving stimulus is caused by differential latencies for moving and flashed stimuli. Vision Research, 40(2), 137–149. 10.1016/S0042-6989(99)00166-2

Yeshurun, Y., & Carrasco, M. (1998). Attention improves or impairs visual performance by enhancing spatial resolution. Nature, 396(6706), Article 6706. 10.1038/23936

Yeshurun, Y., & Carrasco, M. (1999). Spatial attention improves performance in spatial resolution tasks1Parts of this study were presented at the Annual Meeting of the Association for Research in Vision and Ophthalmology (May 1997) and at the Annual Meeting of the Psychonomics Society (November 1997) and published in Abstract format (Yeshurun and Carrasco, 1997and Carrasco and Yeshurun, 1997, respectively).1. Vision Research, 39(2), 293–306. 10.1016/S0042-6989(98)00114-X

Yook, J., Lee, L., Vossel, S., Weidner, R., & Hogendoorn, H. (2022). Motion extrapolation in the flash-lag effect depends on perceived, rather than physical speed. Vision Research, 193, 107978. 10.1016/j.visres.2021.107978

Zhong, S.-h., Ma, Z., Wilson, C., Liu, Y., & Flombaum, J. I. (2014). Why do people appear not to extrapolate trajectories during multiple object tracking? A computational investigation. Journal of Vision, 14(12), 12–12. 10.1167/14.12.12

