## Appendix A for "When visual attention is divided in the flash-lag effect"

**Appendix A: Descriptive statistics of FLE magnitude**

**Table A1.**

Descriptive statistics of FLE magnitudes across attention conditions.

| ID | 1 | | | 2 | | | 3 | | | 4 | | |
|---|---|---|---|---|---|---|---|---|---|---|---|---|
| | Range | Skewness | Kurtosis | Range | Skewness | Kurtosis | Range | Skewness | Kurtosis | Range | Skewness | Kurtosis |
| 1 | -12.4–50.6 | 0.5 | 3.2 | -24.4–58.4 | 0.0 | 2.8 | -33.2–178.4 | 2.8 | 18.5 | -21.2–105.0 | 1.2 | 6.5 |
| 2 | -19.4–53.4 | 0.9 | 4.4 | -24.0–53.6 | 0.5 | 3.9 | -32.4–102.8 | 1.7 | 11.5 | -19.0–29.0 | -0.2 | 3.4 |
| 3 | -16.6–90.8 | 1.0 | 5.4 | -96.0–93.2 | -0.8 | 7.3 | -20.8–78.8 | 0.1 | 2.8 | -60.4–96.8 | 0.1 | 3.7 |
| 4 | -16.2–139.2 | 3.3 | 22.3 | -24.6–90.6 | 1.1 | 7.5 | -44.8–76.4 | 0.3 | 4.8 | -22.2–60.8 | 0.1 | 3.7 |
| 5 | -28.2–121.8 | 1.6 | 10.0 | -49.8–118.8 | 0.9 | 9.0 | -48.4–65.8 | -0.2 | 4.0 | -114.4–62.0 | -1.9 | 14.5 |
| 6 | -21.8–56.6 | 0.4 | 4.5 | -19.0–51.6 | 0.0 | 2.8 | -23.0–54.8 | 0.1 | 2.8 | -28.2–60.2 | -0.1 | 3.6 |
| 7 | -56.4–74.4 | -0.1 | 3.3 | -85.4–121.4 | -0.1 | 4.5 | -131.2–174.2 | -0.2 | 8.8 | -47.6–148.6 | 0.8 | 4.5 |

| | | | | | | | | | | | | |
|---|---|---|---|---|---|---|---|---|---|---|---|---|
| 8 | -19.0–58.0 | 0.9 | 4.7 | -11.0–55.6 | 0.6 | 3.3 | -13.4–51.8 | 0.4 | 2.5 | -24.4–64.8 | 0.5 | 4.8 |
| 9 | -113.0–115.2 | -0.4 | 6.0 | -131.2–172.2 | -0.2 | 5.8 | -90.0–161.2 | 0.5 | 5.7 | -174.6–161.4 | -0.8 | 7.4 |
| 10 | -23.2–50.0 | 0.7 | 3.6 | -28.2–61.6 | 0.7 | 4.8 | -20.4–61.2 | 0.2 | 3.0 | -20.4–67.6 | 0.4 | 4.0 |
| 11 | -172.6–179.0 | -0.5 | 3.2 | -175.8–179.0 | -0.5 | 3.1 | -153.2–174.0 | -0.2 | 2.7 | -163.0–174.6 | -0.3 | 2.6 |
| 12 | -11.2–125.4 | 1.9 | 8.3 | -23.2–100.0 | 0.7 | 4.1 | -73.4–152.6 | 0.7 | 8.0 | -22.6–135.0 | 1.3 | 7.8 |
| 13 | -19.0–53.4 | 0.4 | 3.3 | -18.0–61.0 | 0.4 | 2.8 | -18.0–73.0 | 0.6 | 3.4 | -16.6–62.4 | 0.3 | 2.6 |
| 14 | -23.0–48.0 | 0.3 | 3.6 | -30.0–41.0 | -0.2 | 3.2 | -20.0–52.8 | 0.0 | 3.4 | -47.2–53.0 | -0.3 | 4.6 |
| 15 | -20.8–52.0 | 0.6 | 4.0 | -136.2–56.0 | -3.9 | 31.1 | -22.2–57.6 | 0.4 | 3.4 | -8.8–62.6 | 1.0 | 4.3 |
| 16 | -18.0–57.0 | 0.1 | 3.0 | -25.4–69.4 | 0.2 | 3.3 | -15.8–74.4 | 0.4 | 2.8 | -32.8–67.0 | 0.2 | 3.1 |
| 17 | -161.6–165.4 | -1.1 | 6.8 | -152.8–163.2 | -0.7 | 7.4 | -53.6–178.6 | 1.4 | 6.7 | -139.4–174.6 | -0.3 | 8.3 |
| 18 | -163.0–167.6 | -0.4 | 11.9 | -142.6–171.0 | 0.0 | 10.0 | -39.2–161.2 | 1.3 | 5.6 | -67.4–162.4 | 1.3 | 7.2 |

| | | | | | | | | | | | | |
|---|---|---|---|---|---|---|---|---|---|---|---|---|
| 19 | -40.6–94.0 | 0.8 | 4.5 | -23.2–105.2 | 0.7 | 4.4 | -30.6–158.2 | 1.6 | 8.8 | -175–175.2 | -1.4 | 13.3 |
| 20 | -62.4–60.4 | -0.3 | 3.9 | -43.4–53.4 | -0.3 | 2.9 | -50.8–75.4 | 0.1 | 3.8 | -24.6–53.6 | 0.0 | 2.6 |
| 21 | -35.6–173.6 | 2.4 | 9.8 | -34.6–169.0 | 2.3 | 11.9 | -52.2–153.6 | 1.3 | 8.5 | -27.8–90.6 | 0.6 | 3.4 |
| 22 | -28.2–62.6 | -0.1 | 3.6 | -10.2–60.6 | 0.7 | 3.3 | -31.8–53.8 | -0.3 | 3.1 | -21.8–56.6 | 0.1 | 2.7 |
| 23 | -135.8–54.8 | -2.7 | 20.3 | -46.6–72.6 | 0.0 | 3.4 | -39.2–65.0 | 0.0 | 3.0 | -29.2–65.0 | 0.4 | 3.3 |
| 24 | -132.0–117.4 | -1.2 | 16.4 | -21.2–84.8 | 0.9 | 4.0 | -30.6–155.0 | 2.1 | 13.5 | -27.2–158.0 | 2.1 | 13.0 |

**Table A2.**

One-sample *t*-test (one-tailed) results against zero.

| Bars | *M* | *SD* | $t_{23}$ | *p* |
|---|---|---|---|---|
| 1 | 15.8° | 6.4° | 12.1 | < 0.001 |
| 2 | 19.0° | 9.0° | 10.3 | < 0.001 |
| 3 | 19.4° | 6.5° | 14.6 | < 0.001 |
| 4 | 19.2° | 7.2° | 13.0 | < 0.001 |
