## Appendix B for "When visual attention is divided in the flash-lag effect"

**Appendix B: Descriptive statistics of FLE variability**

**Table B1.**

One-sample *t*-test (one-tailed) results against zero.

| Bars | *M* | *SD* | $t_{23}$ | *p* |
|---|---|---|---|---|
| 1 | 130.2% | 45.3% | 14.1 | < 0.001 |
| 2 | 118.4% | 36.2% | 16.0 | < 0.001 |
| 3 | 114.0% | 32.2% | 17.3 | < 0.001 |
| 4 | 115.4% | 32.2% | 17.6 | < 0.001 |
