## Appendix C for "When visual attention is divided in the flash-lag effect"

### Appendix C: Hemifield effects in the FLE

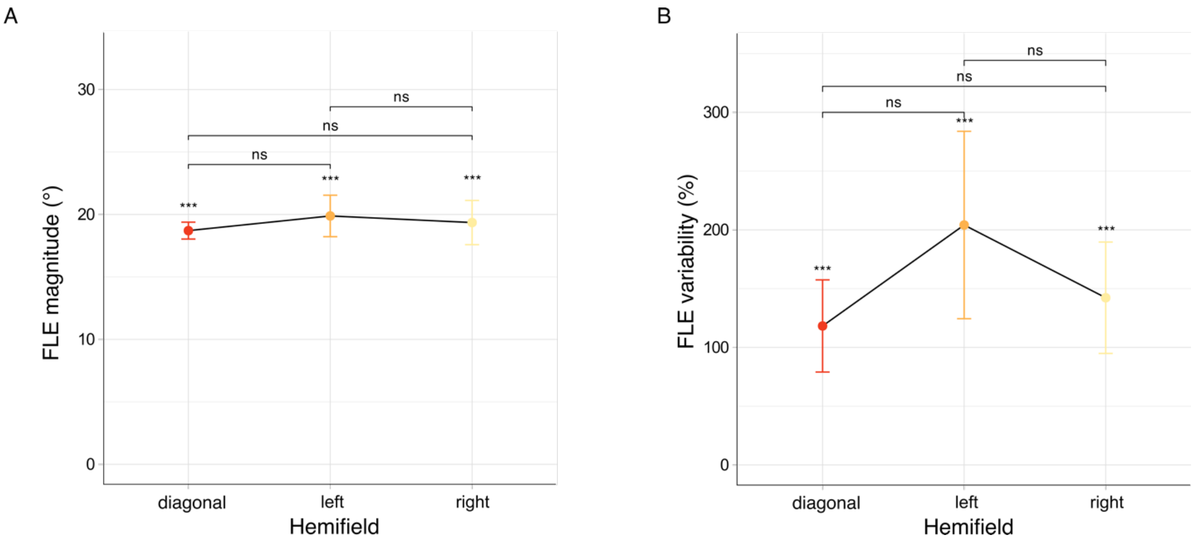


**Figure C1.** Hemifield results. Previous studies using linear motion have demonstrated anisotropies in the left and right sides of the visual field (Kanai et al., 2004; Shi & Nijhawan, 2008; Suzuki et al., 2023). Notably, the FLE tends to be more substantial when stimuli are presented in the left hemifield. Here, we aimed to test whether the effect of divided attention could be pronounced with these anisotropies, particularly within the attend-to-two condition. To achieve this, we examined the magnitude and variability of the FLE on trials where participants were cued to two quadrants within their left, right, or diagonally across the upper and lower visual hemifields. (**A**) The FLE magnitudes are plotted as a function of the to-be-attended hemifields. All values are reported as means ± within-subject standard error of the mean (S.E.M., error bars). ns (not significant) denotes $p > 0.05$. Although the FLE magnitude in the left hemifield was marginally higher than in both the right or diagonal hemifields, no significant modulation of attention was detected (one-way ANOVA, *hemiField:* $F(2,46) = 0.2$, $p = 0.7$). (**B**) The same as described in **A** but for FLE variability (one-way ANOVA, *hemiField:* $F(2,46) = 0.6$, $p = 0.5$).
